# Identification of Iron Deficiency-Induced Chlorosis Tolerant Lines from Wild Rice Germplasm

**DOI:** 10.1101/2023.03.30.534996

**Authors:** Rahul Kumar

## Abstract

The cultivation of transplanted rice has led to depletion of groundwater levels in many parts of the world. Direct-seeded rice under aerobic conditions is an alternative production technology that requires much less water. However, under aerobic conditions, iron in the soil is oxidized from ferric to ferrous, which is not easily available for rice, resulting in iron deficiency-induced chlorosis (IDIC) and a drastic reduction in seed yield. Cultivated rice genotypes have limited variations for IDIC tolerance, therefore, wild *Oryza* germplasm could be a better source for IDIC tolerance. In this study, 313 *Oryza* accessions were evaluated for IDIC tolerance at the tillering stage under aerobic conditions using IDIC rating, SPAD value, and iron content in leaves. Based on these parameters, twenty IDIC tolerant lines were selected. These 20 lines exhibited no chlorosis, high SPAD values, and iron content, while 8 cultivated controls showed mild to high chlorosis symptoms and low SPAD and iron content. In a subsequent year, the selected lines were evaluated again to confirm their tolerance, and they exhibited similar levels of tolerance. These accessions may be useful for developing IDIC-tolerant cultivars for aerobic rice cultivation, and future study to understand the molecular mechanism of IDIC tolerance in rice.

## Introduction

Rice (*Oryza sativa* L., 2n=24) is an important cereal and the primary food source for over one-third of the world’s population (Khush, 2013). It is the second-largest crop grown globally, with 160 million hectares of rice cultivation worldwide (FAOSTAT, 2021). Rice is a staple crop in many countries, and over 90% of its production is consumed in Asia (Khush, 2013). Despite its significance as a food source, rice cultivation poses significant challenges, including high water usage. Irrigated agriculture accounts for 70-85% of global water usage, with rice being one of the most water-consuming crops (Gleick, 2003). This excessive water usage in rice cultivation has resulted in the depletion of groundwater levels in many rice-growing regions worldwide, which could have catastrophic consequences for future global rice production. Hence, the development of a new ideotype of rice that demands less water than the existing cultivation system is very important.

The high-water requirement of rice poses a threat to future rice production, as the groundwater levels in many rice-growing regions are depleting (Tuong et al., 2005). Rice production in many high-yielding regions is at risk due to groundwater depletion, and there is a need to produce more rice with less water to sustain productivity (Liu et al., 2019). Furthermore, the labor-intensive nature of transplanted rice has resulted in many farmers moving away from rice cultivation, necessitating a shift towards more efficient rice cultivation practices (Yadav et al., 2019). Aerobic rice cultivation offers several advantages over traditional rice cultivation methods, such as increased water-use efficiency, reduced greenhouse gas emissions, lower cultivation costs, and decreased labor requirements (Pang et al., 2021). Bouman (2001) suggested that aerobic rice could be grown under irrigated conditions, much like upland crops such as wheat and maize, by cultivating high-yielding rice varieties in direct-sown, non-puddled, aerobic soils with irrigation. This method of cultivation is known as direct-seeded aerobic rice and could potentially serve as an alternative to cope with depleting underground water and labor shortages in rice production (Farooq et al., 2019).

Iron deficiency-induced chlorosis (IDIC) is a major constraint in aerobic rice cultivation, which significantly reduces crop yield and quality (Fan et al., 2012; Shi et al., 2012; Nogiya et al., 2016; Nogiya et al., 2019). Although some studies have been conducted on IDIC in rice, there is still a lack of information on screening genotypes for IDIC under aerobic conditions in rice germplasm (Sperotto et al., 2012). Additionally, cultivated species have limited variation for these traits, making it difficult to develop iron-efficient cultivars. IDIC is a physiological disorder that affects the growth and yield of rice under aerobic conditions. In rice cultivated under aerobic conditions, the solubility of iron in the soil is limited due to its oxidation to the insoluble ferric form (Fan et al., 2012; Shi et al., 2012; Nogiya et al., 2016; Nogiya et al., 2019). IDIC is characterized by the interveinal chlorosis of young leaves, reduced plant growth, and decreased yield (Kumar et al., 2013). The mechanism of IDIC in rice under aerobic conditions involves the regulation of iron uptake, translocation, and utilization (Cakmak, 2008; Sperotto et al., 2012).

Wild rice species possess a vast array of desirable alleles for abiotic stress tolerance, including IDIC tolerance, as they have evolved under diverse environmental conditions (Brar and Khush, 1997). The *Oryza* genus includes 21 wild and two cultivated species of rice, and over 70% of the genetic variation in this genus is attributed to wild species (Tanksley and McCouch, 1997). *O. sativa* (L.), which originated from *O. nivara* and *O. rufipogon*, is grown worldwide, while *O. glaberrima* (Steud.), which originated in West Africa from *O. barthii* (A. Chev.), is grown on a limited scale (Vaughan et al., 2003).

IDIC is a common problem in rice cultivation, particularly under aerobic conditions. However, limited research has been conducted to date on screening rice genotypes for IDIC tolerance under such conditions, and cultivated species exhibit limited variation for these traits (Fan et al., 2012; Shi et al., 2012; Nogiya et al., 2016; Nogiya et al., 2019). Wild rice species have evolved under diverse environmental conditions and harbor desirable alleles for various biotic and abiotic stresses, making them a promising genetic resource for improving IDIC tolerance in cultivated rice. Punjab Agricultural University in Ludhiana, India, has a vast collection of wild rice species procured from the International Rice Research Institute in the Philippines. The purpose of this study was to screen wild *Oryza* species for IDIC tolerance at the tillering stage under direct-seeded aerobic conditions.

## Materials and Methods

### Climate

The experiment was conducted at Punjab Agricultural University, Ludhiana, utilizing both field and laboratory facilities. The study site has geographical coordinates of 30°56’ N latitude and 75°52’ E longitude, and a mean altitude of 247 meters above sea level. The area has a semi-arid sub-tropical climate with distinct seasonal variations. The summer season, which lasts from April to June, is hot and dry, while the monsoon season, from July to September, is hot and humid. The winter season can be divided into two parts, mild winter from October to November and cold winter from December to February. The soil was found to have a Fe content of 4.86 ppm.

### Plant Materials

In this study, a total of 313 rice genotypes were used as plant material, consisting of six accessions of *O. glaberrima*, 105 accessions of *O. rufipogon*, 193 accessions of *O. nivara*, one accession of *O. barthii*, and 8 cultivated *O*. sativa genotypes (Fig. 1). The experiment was conducted in a Randomized Block Design (RBD) with three replications. Paired rows of 1.5m length were sown for each entry with a row spacing of 30 cm under dry direct seeded conditions.

**Figure 1:**
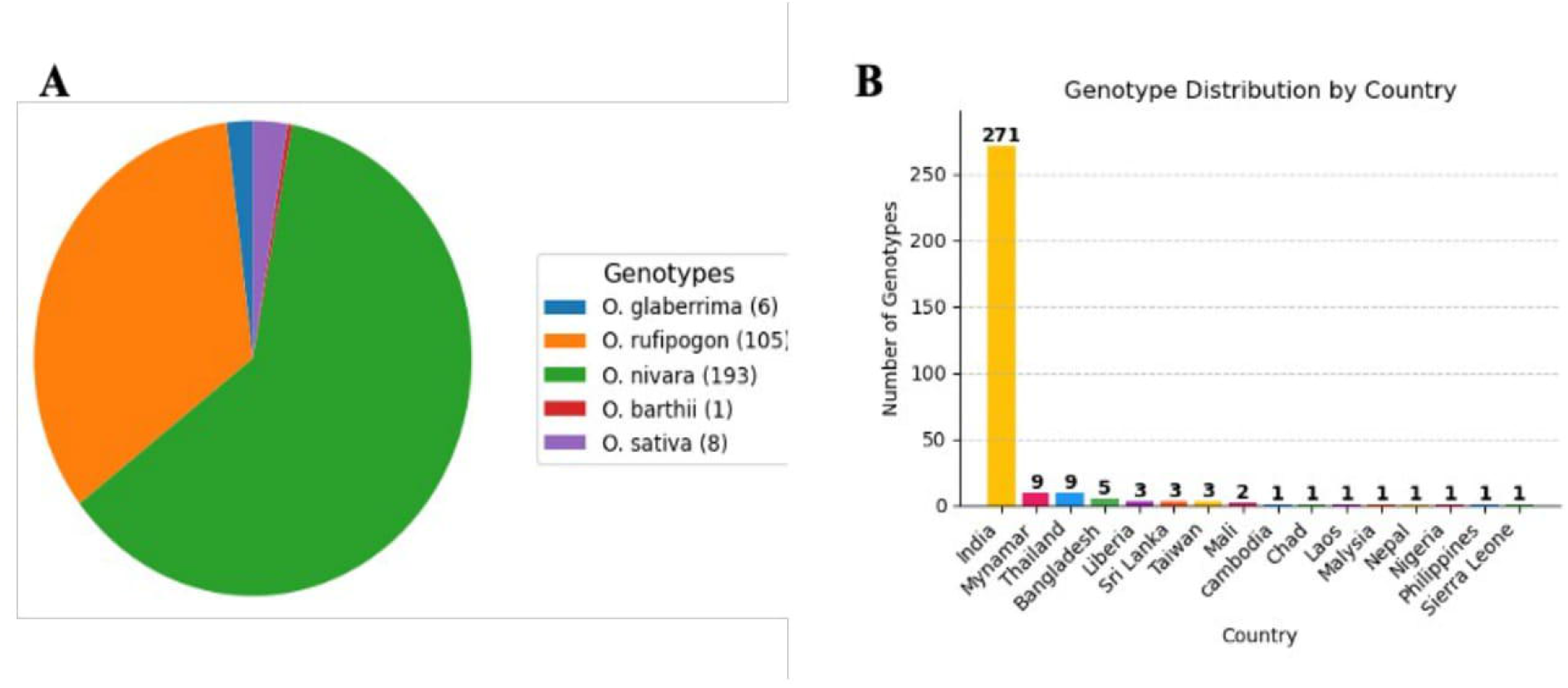
Information of the genotype used for the experiment. **(A)** Species wise distribution of the germplasm into five group. Majority of the genotypes were fall into *O. nivara* and *O. rufipogon*. **(B)** County wise distribution of wild germplasm. Majority of wild germplasm were belonged to India.

### Irrigation applied during experiment

During the rice season, the field was irrigated immediately after sowing and thereafter on a weekly basis, depending on the amount of rainfall, to maintain the required soil moisture levels. In instances where there was rainfall, the scheduled irrigation was skipped, and additional irrigation was administered at 7-day intervals after the water had drained from the field.

### Screening method for IDIC tolerance of wild germplasm

To evaluate IDIC tolerance, the rice genotypes were evaluated on a 1-5 scale after 28 days of sowing. The ratings were based on the degree of chlorosis observed in the plants. A score of 1 indicated normal, green plants without any chlorosis, while a score of 2 indicated slight yellowing of the upper leaves. A score of 3 indicated interveinal chlorosis in the upper leaves without any stunting of growth or necrosis, and a score of 4 indicated interveinal chlorosis of the upper leaves with some stunting of growth or necrosis of plant tissue. A score of 5 indicated severe chlorosis with stunted growth and necrosis of the youngest leaves and growing point (Wang et al., 2008). The ratings were recorded in each plot to determine the IDIC tolerance of the rice genotypes.

### Chlorophyll measurement and leaf area index

The SPAD meter was used to measure chlorophyll content in the leaf. The SPAD value of each entry was determined by after 28 days of sowing using a SPAD meter to measure the middle portion of the index leaf. Five randomly selected plants from each entry were measured to ensure accuracy, while wet leaves and plants that were widely spaced, tall, or short were avoided during the measurement process. Leaf area index (LAI) was recorded at tillering using a digital plant canopy imager (Model CI-110/CI-120, CIDInc, USA).

### Iron content in the leaves

After four weeks of sowing, the plants were uprooted, thoroughly washed twice with deionized water, and then dried in a hot air oven at 65°C for 3 days. The dried leaves were finely ground to analyze the Fe concentration. To analyze the total Fe, the dried samples were digested in a diacid mixture of HNO3 and HClO4 (3:1). The concentrations of Fe were determined using the atomic absorption spectrophotometry method described by Isaac and Kerber (1971).

### Statistical analysis

ANOVA was performed to compare the Fe content in the leaf with SPAD values, using the python. Furthermore, correlation analysis was carried out to investigate the correlation between IDIC with iron content and SPAD reading using the python.

## Results

### Wild *Oryza* germplasm exhibited significant variation for IDIC tolerance under aerobic field conditions

IDIC is a major limiting factor for the adaptation of aerobic rice, affecting plant growth and yield. IDIC was scored on a 1-5 scale at the tillering stage, as described in the Materials and Methods section. A significant variation was observed between the 313 rice genotypes tested. While 20 genotypes remained green throughout the growing season (Rating 1), the remaining wild germplasm lines and conventional cultivars showed mild to severe chlorosis and stunting symptoms (Rating 2-5) (Table 1, Fig. 2, Fig. 3). Among the IDIC-tolerant genotypes, 11 belonged to the *O. nivara* group, and 9 to the *O. rufipogon* group (Table 1). Subsequent year, 20 selected IDIC-tolerant lines and eight cultivars were evaluated for their tolerance to IDIC under aerobic conditions. Similar IDIC responses were observed compared to the previous year (Table 1). Of the 20 wild germplasm lines, 17 showed a rating of 1, while three showed a rating between 1 and 2. In contrast, all cultivars had a rating of 2 or higher, with Feng-ai-zai showing a rating of 5. The correlation analysis showed that IDIC rating was negatively related to SPAD value (correlation coefficient = −0.69), iron content (correlation coefficient = −0.56), and LAI (correlation coefficient = −0.28) (Fig. 4). These findings suggest that there is considerable variation in IDIC tolerance among rice genotypes, with wild germplasm lines showing greater tolerance compared to conventional cultivars.

**Table 1.** List of the 20 selected tolerant and cultivated genotypes with IDIC, SPAD, LAI and iron content. * A statistical difference was observed among the genotypes.

| Species | First year |  |  |  | Second year |  |
| --- | --- | --- | --- | --- | --- | --- |
|  | IDIC rating* | SPAD reading* | Fe content (ppm)* | LAI* | IDIC rating* | SPAD Reading* |
| *O. rufipogon* (IR105491) | 1±0 | 32.5±1.8 | 1294±131 | 1.4±0.13 | 1±0 | 33.1±1.3 |
| *O. rufipogon* (CR100334A) | 1±0 | 31.6±0.8 | 1119±79 | 1.5±0.21 | 1±0 | 30.9±0.9 |
| *O. rufipogon* (CR100334B) | 1±0 | 31.2±1.1 | 1261±113 | 2.1±0.19 | 1±0 | 33.9±1.3 |
| *O. rufipogon* (CR100436) | 1±0 | 32.7±1.9 | 1664±143 | 1.4±0.11 | 1.3±0.47 | 30.2±1.6 |
| *O. rufipogon* (CR100462) | 1±0 | 32.8±2.1 | 1042±92 | 1.4±0.16 | 1±0 | 32.4±1.7 |
| *O. rufipogon* (IR80762A) | 1±0 | 32.9±0.4 | 1229±75 | 1.8±0.09 | 1±0 | 34.3±0.9 |
| *O. rufipogon* (CR100030) | 1±0 | 33.3±1.4 | 2127±193 | 1.2±0.13 | 1±0 | 32.8±1.6 |
| *O. rufipogon* (CR100001) | 1±0 | 35.3±1.3 | 1056±69 | 2.2±0.10 | 1.3±0.47 | 34.6±1.6 |
| *O. rufipogon* (CR100484) | 1±0 | 32.5±1 | 1085±82 | 1.4±0.09 | 1±0 | 33.1±1.7 |
| *O. nivara* (CR100454) | 1±0 | 31±0.6 | 1005±63 | 1.1±0.14 | 1±0 | 31.8±1.6 |
| *O. nivara* (CR100363) | 1±0 | 32.3±2.2 | 2101±173 | 1.4±0.12 | 1±0 | 31.2±1.7 |
| *O. nivara* (CR100371) | 1±0 | 33.8±1.3 | 1055±103 | 1.1±0.12 | 1±0 | 33.2±1.1 |
| *O. nivara* (CR100003) | 1±0 | 36±1.6 | 1802±144 | 1.5±0.16 | 1±0 | 34.7±1.9 |
| *O. nivara* (IR82018) | 1±0 | 29.7±2.5 | 1287±103 | 1.6±0.17 | 1.66±0.47 | 31.9±1.9 |
| *O. nivara* (CR100124A) | 1±0 | 29.8±0.8 | 1081±89 | 1.5±0.10 | 1±0 | 31.5±0.9 |
| *O. nivara* (CR100127) | 1±0 | 30.3±1.4 | 1662±123 | 1.6±0.12 | 1±0 | 29.8±1.1 |
| *O. nivara* (CR100337) | 1±0 | 33.3±1.1 | 1299±91 | 1.3±0.12 | 1±0 | 33.9±0.8 |
| *O. nivara* (CR100426) | 1±0 | 31.6±1.9 | 1079±59 | 1.2±0.08 | 1±0 | 32.3±1.7 |
| *O. nivara* (CR100360) | 1±0 | 33.5±0.9 | 1508±179 | 1±0.10 | 1±0 | 31.5±1.1 |
| *O. nivara* (CR100429) | 1±0 | 29.4±1.6 | 1318±132 | 1.3±0.07 | 1±0 | 30.1±1.5 |
| PAU 201 | 2.7±0.27 | 27.9±1.2 | 565±51 | 1.7±0.11 | 3±0 | 31.7±0.9 |
| Rasi | 4±0 | 29.2±1.7 | 560±60 | 1.2±0.09 | 3.3±0.27 | 29.7±1.3 |
| Basmati 370 | 1.67±0.47 | 23.9±0.9 | 580±44 | 1.2±0.07 | 2±0.0 | 27.5±2.1 |
| Cheng-Hui 448 | 4±0 | 26.9±2.1 | 682±77 | 1.2±0.10 | 4±0 | 26.9±1.1 |
| Feng-ai-zai | 5±0 | 28.2±2.8 | 847±92 | 1.1±0.11 | 5±0 | 29.1±1.9 |
| IR 64 | 3.3±0.27 | 28.3±1.7 | 734±55 | 1.2±0.10 | 4±0 | 26.8±1.9 |
| Lemont | 1.67±0.47 | 31.1±0.8 | 851±73 | 1.6±0.08 | 2±0.0 | 32.1±1.8 |
| Nagina 22 | 3±0 | 23.6±1.9 | 765±61 | 0.9±0.08 | 3.3±0.27 | 26.1±1.4 |

**Figure 2:**
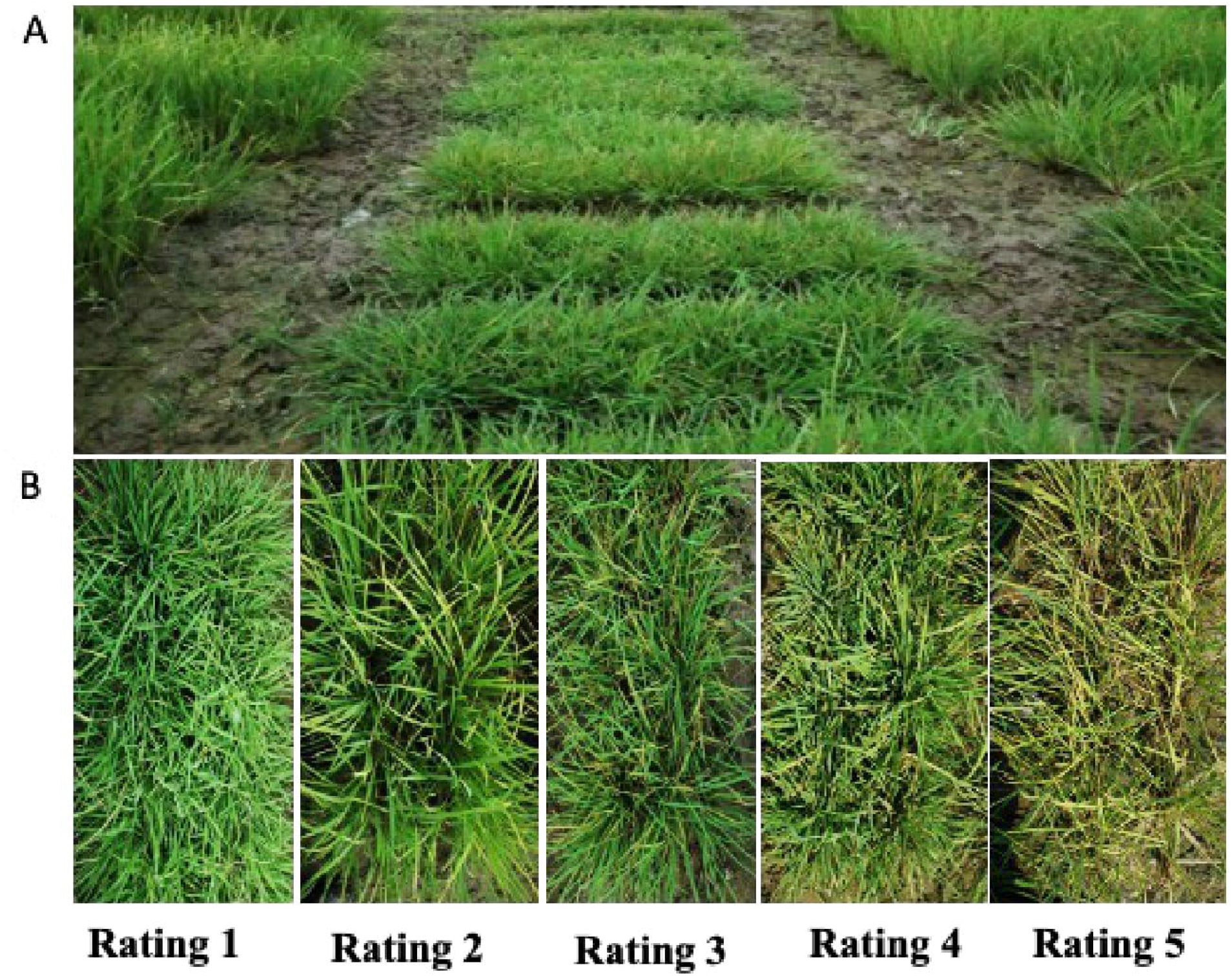
Evaluation of wild *Oryza* germplasm for iron deficiency induced chlorosis tolerance under aerobic conditions. **(A)** Wild *Otyza* wild germplasm comprise of 313 genotypes along with cultivars were evaluated for iron deficiency induced chlorosis tolerance under aerobic conditions. IDIC rating was taken after 28 days after sowing. **(B)** A 1-5 rating was given to score iron deficiency induced chlorosis tolerance. A score of 1 indicated no chlorosis and normal, green plants, while a score of 2 indicated slight yellowing of the upper leaves. A score of 3 indicated interveinal chlorosis in the upper leaves, but no obvious stunting of growth or death of leaf tissue (necrosis), and a score of 4 indicated interveinal chlorosis of the upper leaves with some stunting of growth or necrosis of plant tissue. A score of 5 indicated severe chlorosis with stunted growth and necrosis of the youngest leaves and growing point.

**Figure 3:**
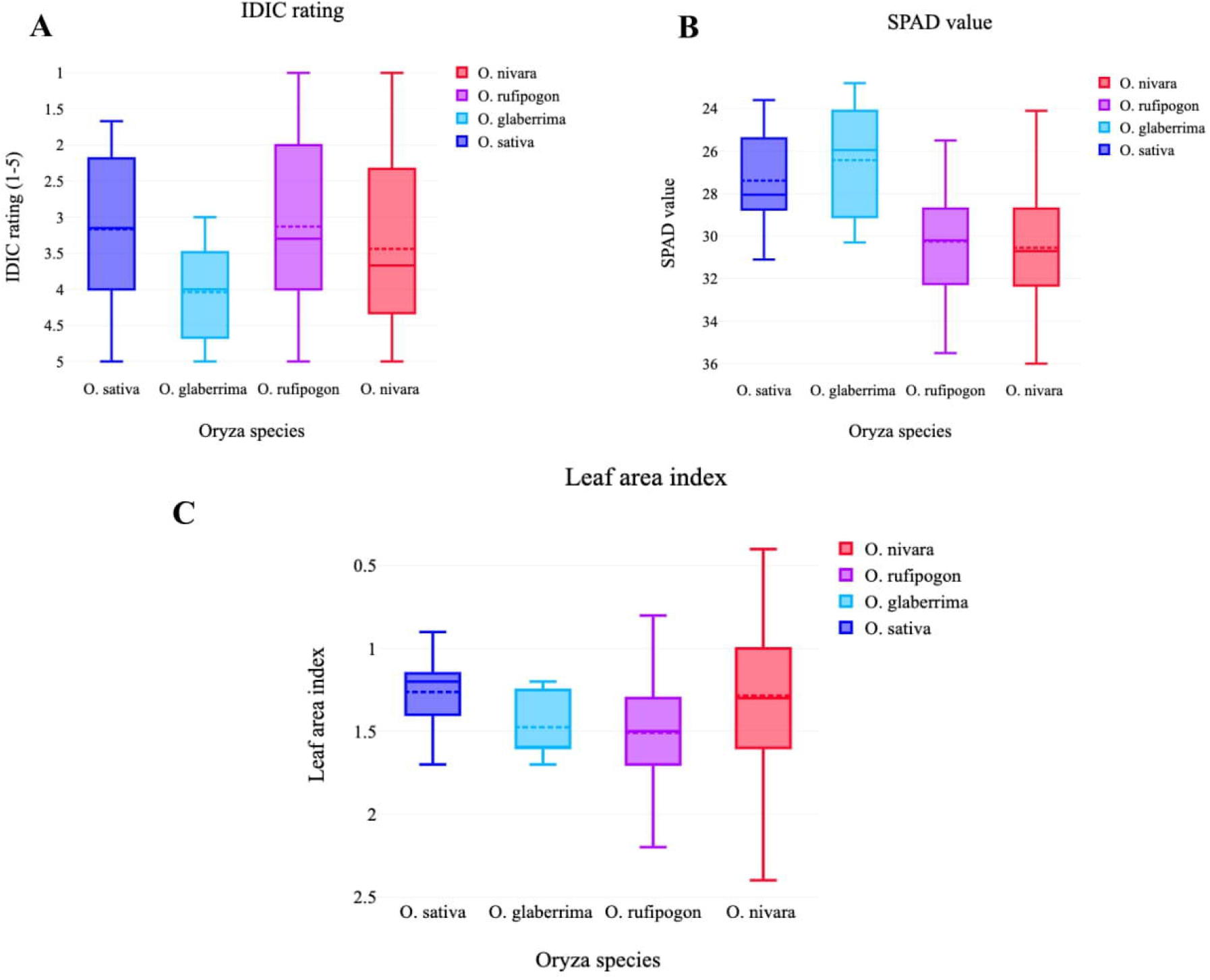
Characterization of wild germplasm for IDIC tolerance at the tillering stage. **(A)** A 1-5 rating was given to score iron deficiency induced chlorosis tolerance. A score of 1 indicated no chlorosis and normal, green plants, while a score of more than 1 indicated susceptibility to IDIC. **(B)** SPAD value (chlorophyll content) of the germplasm at the tillering stage. **(C)** LAI of the germplasm at the tillering stage.

**Figure 4:**
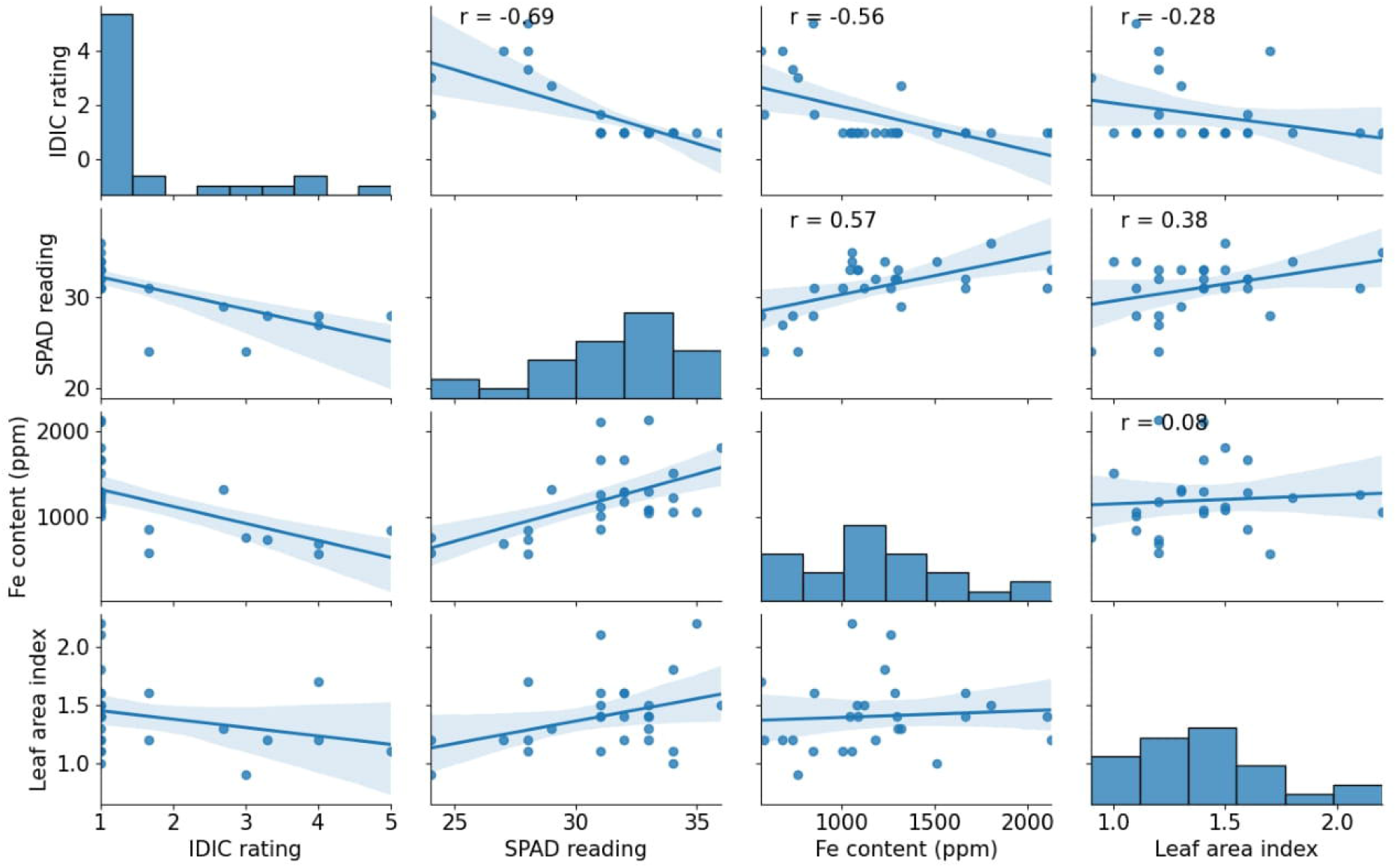
Correlation analysis between IDIC raring, SPAD value, LAI and iron content. A strong correlation was observed between IDIC rating and iron content, while moderate correlation was observed IDIC with SPAD value and LAI.

### Wild *Oryza* germplasm exhibited significant variation for SPAD value

The SPAD meter was used to measure chlorophyll content and relative greenness of leaves in the different genotypes. The results showed significant variations in the SPAD values across the genotypes, with *O. nivara* and *O. rufipogon* species having the highest average SPAD values of 30.8 and 30.6, respectively, followed by *O. sativa* at 28.3 (Table 2). Among the wild genotypes, *O. nivara* (CR 100003) had the highest SPAD value of 36, followed by *O. rufipogon* (CR 100001), *O. nivara* (CR 100371), and *O. nivara* (CR 100360) with values of 35.3, 33.8, and 33.5, respectively (Table 2, Fig. 3). Among the cultivars, Lemont had the highest SPAD value of 31.1, followed by Rasi, IR 64, and Feng-ai-zai with values of 29.2, 28.3, and 28.2, respectively. Among the wild genotype *O. nivara* (CR 100429) had the lowest SPAD value of 29.4, while Nagina 22 had the lowest value of 23.6 in the conventional cultivar group (Table 1). Subsequent year, expectedly IDIC tolerant wild *Oryza* genotypes had higher SPAD value than susceptible cultivated varieties. The correlation analysis showed that SPAD value was negatively related to IDIC rating (correlation coefficient = −0.69) and positively with iron content (correlation coefficient = 0.57), and LAI (correlation coefficient = 0.38) (Fig.4). These results suggest that selected 20 IDIC tolerant genotypes of *O. nivara* and *O. rufipogon* may have a better ability to maintain chlorophyll content and plant health under iron-deficient conditions.

**Table 2.** Mean and range for SPAD value and LAI of 313 genotypes taken after 28 days of sowing.

|  | SPAD value |  |  | LAI |  |  |
| --- | --- | --- | --- | --- | --- | --- |
| Species | Mean | Maximum | Minimum | Mean | Maximum | Minimum |
| *O. nivara* | 30.6 | 36.0<br>(CR100003) | 22.2<br>(CR100123) | 1.3 | 2.6<br>(CR 100007) | 0.4<br>(CR 100399) |
| *O. rufipogon* | 30.8 | 35.3<br>(CR100001) | 25.5<br>(CR100461) | 1.5 | 2.2<br>(CR 100001) | 0.8<br>(CR 100465B) |
| *O. glaberrima* | 26.4 | 30.3<br>(IR102980) | 22.8<br>(IR102489) | 1.5 | 1.7<br>(IR 102489) | 1.2<br>(IR 102980) |
| *O. barthii* | 24.9 | 24.9<br>(IR103580) | 24.9<br>(IR103580) | 1.4 | 1.4<br>(IR103580) | 1.4<br>(IR103580) |
| *O. sativa* | 28.3 | 31.1<br>(Lemont) | 23.6<br>(Nagina 22) | 1.2 | 1.7<br>(PAU 201) | 0.9<br>(Nagina 22) |

### IDIC tolerant lines had more iron content in the leaves than susceptible cultivars

Iron content in the leaves of 20 IDIC tolerant lines and 8 conventional cultivars was determined four weeks days after sowing, and the results showed significant differences between genotypes. The tolerant lines had higher iron content compared to conventional cultivars, with wild *Oryza* germplasm exhibiting a wider range of iron content (1005 to 2121 ppm) compared to conventional cultivars (560 to 847 ppm) (Table 1, Fig. 3). *O. rufipogon* (CR 100030) had the highest iron content of 2127 ppm, followed by *O. nivara* (CR100363), *O. nivara* (CR 100003), and *O. rufipogon* (CR 100436) with iron content of 2101, 1802, and 1664 ppm, respectively (Table 1). Conversely, the highest iron content in conventional cultivars was observed in the Loment cultivar with 851 ppm, followed by Feng-ai-zai, Nagina 22, and IR 64 with iron content of 847, 765, and 734 ppm, respectively. The correlation analysis showed that iron content was negatively related to IDIC rating (correlation coefficient = −0.56) and positively with SPAD value (correlation coefficient = 0.57), and LAI (correlation coefficient = 0.08) (Fig. 4). These findings demonstrate that the IDIC tolerant lines have significantly higher iron content compared to conventional cultivars and that wild *Oryza* germplasm may serve as a valuable genetic resource for aerobic rice.

### Wild germplasm exhibited significant variation for LAI

Significant differences were observed between the genotypes for LAI under aerobic conditions, with average values ranging from 0.9 to 2.6 (Table 2). Among the wild genotypes, *O. rufipogon* (CR 100001) had the highest LAI 2.2, followed by *O. rufipogon* (CR 100334B), *O. rufipogon* (IR 80762A), and *O. nivara* (IR 82018) with values of 2.1, 1.8, and 1.8, respectively (Table 1). In the cultivar genotypes, PAU 201 had the highest LAI with a value of 1.7, followed by Lemont with a value of 1.6. The wild genotype *O. nivara* (CR 100360) had the lowest LAI of 1, while Nagina 22 had the lowest LAI of 0.9 among the conventional cultivar group (Table 1). The correlation analysis showed that LAI was negatively related to IDIC rating (correlation coefficient = −0.28) and positively with SPAD value (correlation coefficient = 0.38), and iron content (correlation coefficient = 0.08) (Fig. 4). These results suggest that there are significant differences in the LAI between the genotypes, with some wild germplasm lines showing higher LAI values compared to conventional cultivars.

## Discussion

Iron deficiency is one of the major constraints in the adaptation of aerobic rice, affecting plant growth and yield. In the present study, a significant genetic variation was observed for IDIC tolerance across *Oryza* the wild germplasm. Our results indicate that wild rice germplasm, particularly *O. nivara* and *O. rufipogon*, have a higher IDIC tolerance than cultivated rice varieties. A moderate to high correlation was observed between IDIC, SPAD value, and iron content. The use of wild rice germplasm in breeding programs may lead to the development of IDIC tolerant rice varieties with higher yield potential and nutritional quality.

In our study, all *Oryza sativa* cultivars showed IDIC under aerobic conditions suggesting lack of genes which can uptake Fe3+ form of iron. The domestication of lowland rice cultivars involved selecting plants that were well-suited for the lowland waterlogged environment, resulting in the loss of many alleles that were important for aerobic rice cultivation (Fuller et al., 2010). As a result of distinct adaptations to their respective environments, lowland and aerobic rice have differences in nutrient availability and uptake (Fan et al., 2012; Shi et al., 2012; Nogiya et al., 2016; Nogiya et al., 2019). The availability of Fe (iron) and Zn (zinc) can become limited under aerobic conditions due to their oxidation (Kirk et al., 20008; Wang et al., 2017). Under anaerobic conditions, iron is available in its ferrous form, which is easily taken up by high-yielding cultivars developed for lowland cultivation (Briat and Lobréaux, 1997). However, under aerobic conditions, iron is insoluble and not available for uptake by rice. In response to iron deficiency stress, plants release iron chelating substances called mugineic acid family phytosiderophores (Ishimaru et al., 2006). These phytosiderophores solubilize inorganic Fe3+ compounds by chelation, and Fe3+-MAs complexes are taken up through a specific transport system in the root plasma membrane (Marschner et al., 1986). Variations in MA synthesizing and iron transporter genes may play a role in IDIC tolerance in resistance lines. Wild species, which have evolved under varying environmental conditions, may possess desirable alleles for aerobic cultivation.

Wild rice germplasm has been extensively demonstrated as a source of stress resistance genes for rice breeding programs (Brar and Khush, 1997). Khan et al. (2020) demonstrated that the introduction of wild rice alleles into cultivated rice significantly improved yield under drought conditions. Similarly, Peng et al. (2020) found that some wild rice species possess genes that confer resistance to bacterial blight. Another study by Feng et al. (2019) characterized the genetic diversity of wild rice germplasm and identified several accessions with high levels of stress tolerance. Li et al. (2019) also showed that wild rice germplasm also contains genes related to cold tolerance. Furthermore, several studies have also shown that wild rice germplasm possesses a high level of genetic diversity, which is essential for the development of new rice cultivars with enhanced stress tolerance (Chen et al., 2019; Li et al., 2019). Similar to these studies, a significant variation for IDIC tolerance was observed in wild germplasm in our study. Identification of twenty IDIC tolerant lines in our study suggests the presence of allelic variations in the phytosiderophores synthesis, and iron transporter genes which potentially can be transferred to high yielding lowland cultivars for aerobic cultivation.

The iron content in IDIC tolerant lines and cultivated susceptible rice varieties was estimated, and results showed significant differences in iron content among genotypes. IDIC tolerant wild genotypes had higher iron content compared to cultivated susceptible rice varieties, with *O. rufipogon* and *O. nivara* having the highest iron content among the wild rice germplasms. Moderately IDIC tolerant Lemont cultivar had the highest iron content among the cultivated varieties which congruent with SPAD value. High level of iron has been reported been reported with IDIC in rice (Kumar et al., 2013). A significant variation in iron content was evident among the twenty IDIC-tolerant genotypes. It is possible that IDIC tolerance in selected lines may be controlled by different mechanisms, as wild germplasm is known to contain a rich diversity of genetic variations. This could be due to the fact that different lines have adapted to iron-deficient conditions through various strategies, such as changes in root architecture, production of iron-chelating compounds, or increased expression of genes involved in iron metabolism. Therefore, the mechanisms underlying IDIC tolerance in different lines may vary depending on the specific genetic variations that have been selected for in each line. In addition to IDIC, we also measured SPAD values as an indicator of chlorophyll content and plant health. Our results showed significant differences in SPAD values among genotypes, with *O. nivara* and *O. rufipogon* having higher SPAD values than *O. glaberrima, O. barthii*, and *O. sativa*. The higher chlorophyll content in wild rice germplasm may be attributed to their ability to adapt to harsh environmental conditions, including iron-deficient soils. it is important to take into account that cultivated varieties typically have higher levels of chlorophyll than wild genotypes under normal growth conditions. Therefore, any differences in SPAD values between wild and cultivated genotypes may not entirely indicate their tolerance to IDIC. The LAI of a plant is an important factor that determines its photosynthetic capacity and yield potential. Our results showed significant differences in LAI among genotypes, with *O. rufipogon* having the highest LAI among the wild rice germplasm and PAU 201 having the highest LAI among the cultivated varieties. IDIC tolerance might contribute to high LAI, but there were genotypic differences for LAI even under normal conditions (data not given), suggesting that LAI might not be good indicator of IDIC tolerance.

In conclusion, our study provides important insights into the genetic variation of IDIC tolerance, chlorophyll content, LAI, and iron content among rice genotypes. Our results indicate that wild rice germplasm, particularly *O. nivara* and *O. rufipogon*, have a higher tolerance to IDIC and higher chlorophyll content, LAI, and iron content than cultivated rice varieties. The use of wild rice germplasm in breeding programs may lead to the development of IDIC tolerant rice varieties with higher yield potential and nutritional quality.

## Author Contributions

RK performed the experiments, data analysis, manuscript writing and editing.

